# Interpretation of Predictions in Drug-Gut Bacteria Interactions Using Machine Learning

**DOI:** 10.1101/2023.03.21.533683

**Authors:** Himanshu Joshi, Meher K Prakash

## Abstract

Gut bacteria play a crucial role in host’s metabolism. Both antibiotic and non-antibiotic drugs affect the gut bacteria ecosystem, which negatively affects the host’s health. Also, gut bacteria metabolize drugs, making them ineffective to the target. The structure-activity relationship studies of drugs have the scope to make them more effective, efficient, and specific to the target. Previous machine learning studies use the available data to predict the activity of drugs and gut bacteria on each other, but these models lack interpretability. Herein, we study the drug-gut bacteria interaction using interpretable machine learning models. In this study, we identify the most important physicochemical features of the drug, which decide the drug-gut bacteria interactions with each other. One of the key findings of this work is that the higher-positive charged drug molecules can inhibit the growth of gut bacteria more. In contrast, the higher-negative charged drug molecules have higher possibility to get metabolized by gut bacteria.

## Introduction

Microbiota present in the gut, sharing a symbiotic relationship with humans, ^1–3^ get killed not only by antibiotic drugs but also by non-antibiotic drugs.^4,5^ Hence, the disturbed gut bacterial population harms the host health.^2,6,7^ Recent in vitro studies show that 27 percent of the non-antibiotic drugs also inhibit the growth of gut bacterial strains.^4,8^ If bacteria get killed by a non-antibiotic drug, which also increases the chance of becoming bacteria antibiotic resistant.^4^ Antibiotic resistance is a massive challenge in drug discovery,^9,10^ and in the future, it may have profound effects.^11–13^ As drugs (antibiotic and non-antibiotic) have adverse effects on gut bacteria, drugs also get metabolized and accumulated by the gut bacteria,^14,15^ which again makes drugs ineffective to the target.^16,17^

In general, drug discovery has two main objectives: first, the drug should efficiently work against pathogens, and second, the drug should have minimal side effects on the hosts’ body.^18–20^ Metabolism of drugs by gut bacteria decreases its’ efficiency to the target,^21^ and if the drug kills gut bacteria, causing dysbiosis, it affects the host body adversely.^22^ This negative activity of drugs and gut bacteria on each other is one of the challenges in drug discovery. Around 90 percent of drug candidates fail in trials, making drug discovery intensive in time and money.^10,23,24^ Previous studies show that drug discovery can be made more efficiently by including machine learning methods in the process.^25^ It can be done by screening the drug candidate before the clinical trials and by predicting computationally new drug molecules, making drug discovery less costly in time and money both.^26–29^

This work focuses on studying two-way interactions of drug-gut bacteria on each other with the help of the physicochemical properties of the drug molecules, as shown in Figure 1 (A). The physicochemical features of drugs mean their physical and chemical properties that affect their behavior and decide the activity drug towards the target.^30,31^ Studying these physicochemical features is essential for drug discovery and predicting drug behavior toward gut bacteria. Computational studies on available data focus on the prediction models but consider these aspects independently. ^17,32^ However, all these studies look at these aspects independently and need more interpretability.

**Figure 1:**
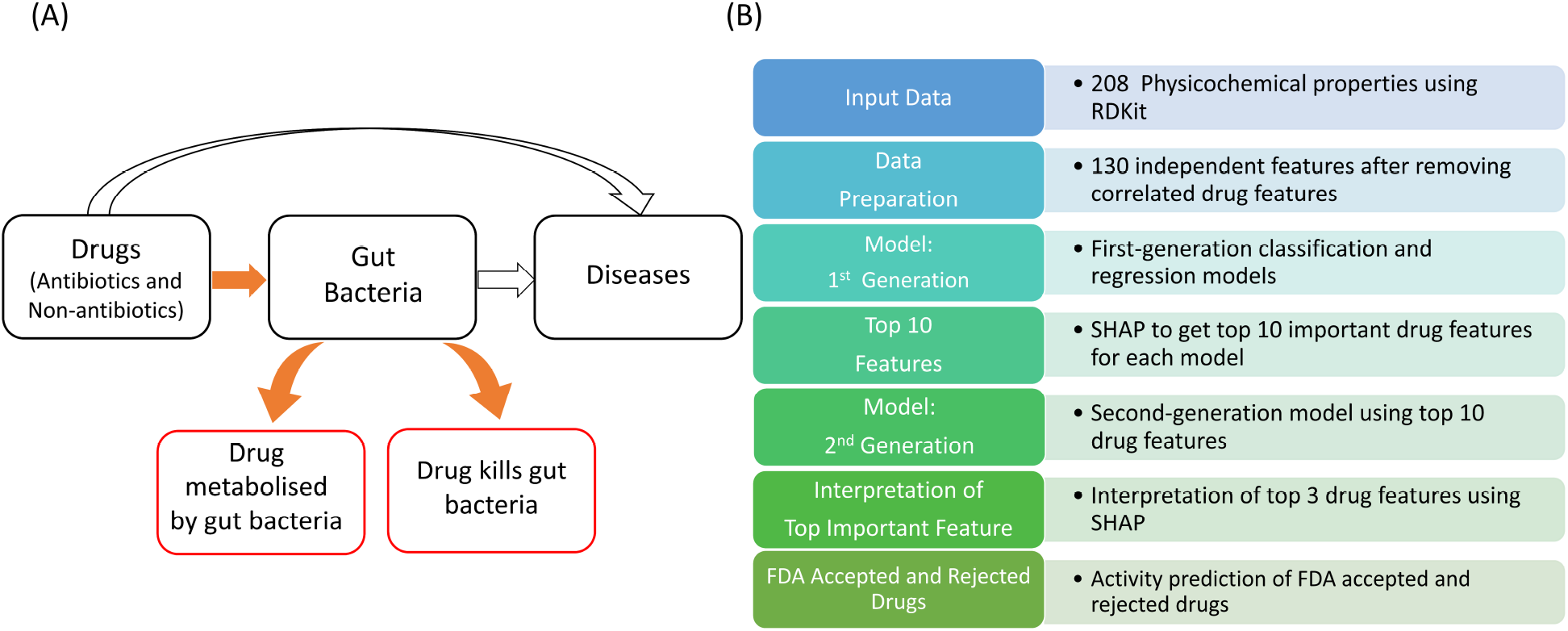
Availability of data and flow diagram of the model. The available data shows that drug (both antibiotic and non-antibiotic) inhibit the growth of gut bacteria (activity data), and other study shows that gut bacteria also metabolize drugs (metabolism data) (A). After removing highly correlated features of the drug molecules, we used a series of two models (XGBoost classification/regression followed by SHAP) to get the top 3 key features from 130 physicochemical properties of drug molecules (B).

So far, there is no interpretable machine learning model for a structure-activity relationship study considering drug metabolism by gut bacteria and drug activity on gut bacteria. Our study identifies which physicochemical features of the drug make it more metabolized by gut bacteria and which make it more harmful to gut bacteria. We found that the maximum partial charge present on the drug molecules is one of the important features in deciding the activity of drug-gut bacteria on each other. One key finding of this study is that cationic drug molecules are more likely to inhibit the growth of gut bacteria, whereas anionic drug molecules are more likely to metabolize more by gut bacteria. Using machine learning to understand the drug-gut bacteria activity on each other using the physicochemical properties of drugs has the potential to make drug discovery more effective to target.

## Methods

### Input Data

We used previously published experimental data of drug activity on gut bacteria (activity data)^4^ and the metabolism of drugs by gut bacteria (metabolism data)^14^ for developing machine learning models. The activity data consisted of P-values with information on drug activity on gut bacteria. We then classified it into two categories: first, if P-value *<* 0.05 (drugs kill the gut bacteria, category 0), and second, if P-value *≥* 0.05 (drugs do not kill the gut bacteria, category 1). The metabolism data contains the percent consumption of drugs by gut bacteria in 24 hours for 20,596 drug-bacteria pairs (271 drugs and 76 gut-bacterial strains). The input data for activity and metabolism are shown in Supporting Information Figure 1 (A) and (B).

### Feature selection

We used various physicochemical properties of the drug molecules as input features for the machine learning model. The physicochemical properties of the drug molecules were generated using smile codes from the drug bank website^33^ and RDkit (a python-based program).^34^ Several of the 208 drug descriptors generated using RDKit were strongly correlated with each other. We used Pearson correlation to remove one out of two correlated features if the value of correlation is more than absolute 0.75. Removing the correlated features, we got 130 independent drug features which we used further in the study. Supporting Information Figure 2 (A) and (B) show the drug features with strong correlation and without correlation, respectively.

### Machine learning model

We used extreme gradient boosting technique (XGBoost)^35^ based regression and classification model for the metabolism and activity data, respectively. To interpret the classification and regression model predictions, we used the SHapley Additive exPlanations (SHAP) algorithm.^36^ For activity data, having two categories, we used XGBoost classification to make 40 prediction models for each bacterium. For metabolism data, we first developed 76 XGBoost regression models and used the criterion used in the parent paper^14^ to classify the drug as metabolized by the bacteria or not. According to this criterion, a drug is considered metabolized if its consumption by bacteria is 20 percent or more in 24 hours (category 0); otherwise, the drug is classified as not getting metabolized (category 1). For both data models, the quality of classification and regression models was assessed based on the metrics: accuracy, precision, recall, and f1-score. We also performed the area under the receiver operating characteristic (AUROC) analysis for activity data.

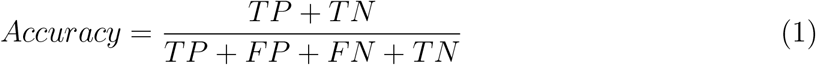

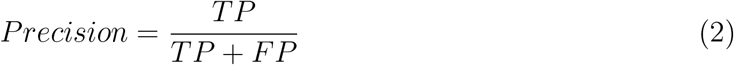

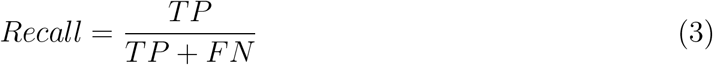

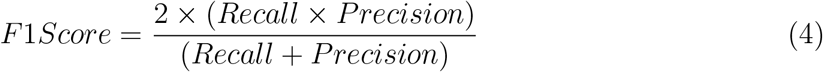

where, TN= True Negative, TP= True Positive, FN= False Negative and FP= False Positive.

### Hyperparameter Tuning

For both the classification and regression models, we optimized the hyperparameters learning rate, max depth, min child width, gamma, and colsample bytree, using the randomised searchCV function of the python module sklearn. ^37^ The ranges of these parameters used for the search are the following; learning rate: 0.05 to 0.3 in increment of 0.05, max depth : 3 to 12 in increment of 1, min child weight: 1 to 7 in increment of 2, gamma : 0.1 to 0.4 in increment of 0.1, colsample bytree : 0.3 to 0.7 in increment of 0.1. The hyperparameters that yielded the best-performing model were used further in the study, the performance assessed with the mean of the AUROC for activity models and accuracy for the metabolism models for the validation sets in a 5-fold cross-validation analysis. ^38^

### Training and Test sets

Both the activity and metabolism data sets are imbalanced, with only 20 percent of the drugs killing the gut bacteria in activity data, and with the higher frequency of the drug not getting metabolized by gut bacteria as shown in the Supporting Information Figure 1 (A) and (B). So, for the activity data, the drugs that do not kill the gut bacteria were downsampled to match the number of drugs that kill bacteria and balance the input activity data. Similarly, the drugs not metabolized by gut bacteria were downsampled to balance the data. The balanced activity and metabolism data is then divided into training (75 percent of the data) and testing (25 percent of the data) to use in the machine learning models.

## Results and Discussion

We developed independent XGBoost models to predict the activity of drugs on gut bacteria (classification models) and drug consumption by gut bacteria (regression models) for each bacterial strain using 130 physicochemical properties of the drugs. We build prediction models with 130 descriptors (first-generation models) to identify the top 10 important descriptors by applying the method of SHAP. Further, we build prediction models with the top 10 descriptors (second-generation models) to identify the top 3 important descriptors by applying the method of SHAP. With this approach, we select the three most important physicochemical properties of drugs out of 130 and interpret the contributions of these features in the models. The flow diagram for the model and analysis is shown in Figure 1 (B).

### Prediction quality of models

The quality of predictions for the first-generation models quantified using various metrics is summarized in Supporting Information Figures 3 (A) and (B) for activity and metabolism data, respectively. For metabolic data, the regression results were used to classify drugs into two categories: first, drug metabolized by gut bacteria if the percent metabolism is more than 20, and second, drug does not get metabolized by gut bacteria if the percent metabolism of the drug is less than 20. The prediction quality for the metabolism data was then evaluated using the classified prediction data. Overall the models performed well in predicting the activity of the drugs (Mean accuracy= 0.84, Standard deviation=0.05) and drug consumption by bacteria (Mean accuracy=0.68, Standard deviation=0.11), and the prediction qualities are comparable with the previous studies.^17,32^ The SHAP algorithm is used to get the top 10 key features of the drugs if the activity data prediction models’ accuracy is more than 0.75 and the metabolism data models’ accuracy is greater than 0.65. Further, we built second-generation models for activity and metabolism data using the top 10 most important features obtained from the first-generation model for 18 strains of bacteria. Figures 2 (A) and (B) show the mean and standard deviation of the metrics used to check the quality of prediction for activity and metabolism data for second-generation models. Supporting Information Figure 4 and 5 shows the prediction qualities of the individual bacterium for the activity and metabolism data, respectively. The prediction qualities of second-generation models are also comparable to the qualities of the previous prediction models.^17,32^

**Figure 2:**
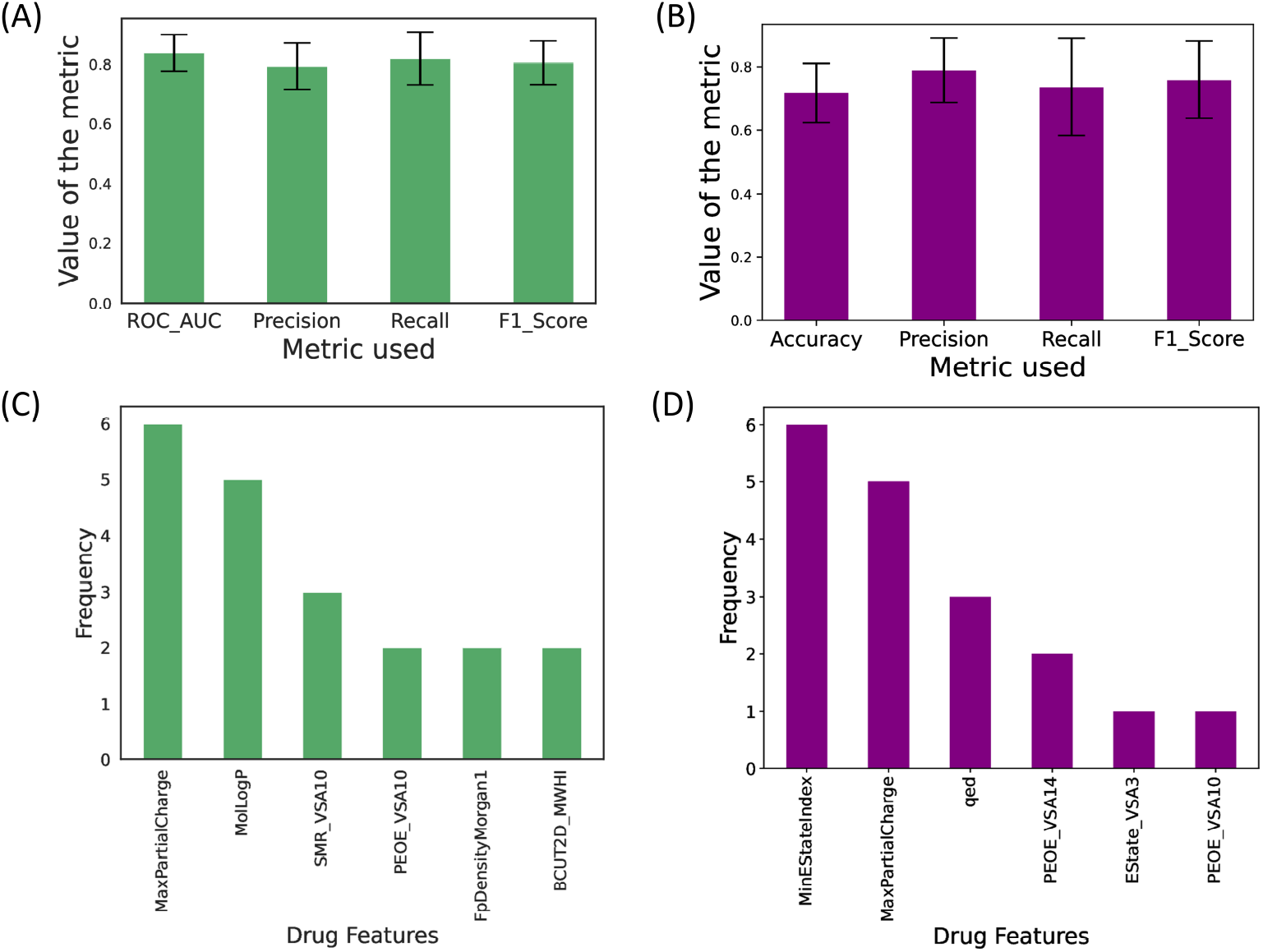
Prediction quality and key drug descriptors in second-generation models: Figure (A) and (B) shows the mean and standard deviation of different metrics used to check the prediction qualities of the individual bacterium model for activity data and metabolism data, respectively. Supporting Information Figure 4 and 5 shows the prediction qualities of the individual bacterium for the activity data and metabolism data, respectively. We used the SHAP algorithm to identify the top key drug physicochemical properties in the activity and prediction data models and plotted the frequency of the key features as shown in (C) and (D) in activity and metabolism data, respectively. The prediction quality and top 10 drug features in first-generation models are given in Supporting Information Figure 3.

### Identification of important features of drugs

To identify the key drug features in first-generation models for each 40 bacterium models for activity data and 74 models for metabolism data, we used the SHAP algorithm, which calculates the contribution of each feature to individual predictions, called the SHAP values. The important features are then identified based on the mean SHAP contribution of the feature to the model. The top ten features from models for each bacteria were selected. While most of the features had a frequency of 1, many were in the top ten features of at least five bacterial strains, as shown in Supporting Information Figure 3 (C) and (D) for the activity and metabolism data, respectively. We followed a similar method as the first-generation model to identify the top drug features in the second-generation prediction model. However, in this case, the top 3 features of the drug from individual models were used. The frequency of different features is shown in Figure 2 (C) and (D) in both the activity and metabolism data models, respectively. The *MaxPartialCharge*, the maximum partial charge value present on any drug molecule, is a common high-frequency drug feature in both activity data and metabolism data but other important features are found to be different for the two data set models. A detailed definition of all these features is given in the Supporting Information.

### Interpretation of key drug features

#### Contribution of drug features to the models

We used the shap.summary.plot of the SHAP algorithm^36^ to interpret the contribution of key physicochemical properties of drugs in the activity and metabolism data models. A negative SHAP value means the drug has a negative effect on gut bacteria, and a positive value of SHAP means the drug will not have a negative effect on gut bacteria.Figure 3 (A) and (C) show the summary of the contribution of important drug features. Higher values of PEOE VSA10, *MaxPartialCharge*, SMR VSA10, and MLogP for any drug give negative SHAP values means these features make the drug more harmful to gut bacteria as shown in figure 3 (A) and smaller values of *MaxPartialCharge*, SlogP VSA3 for any drug give negative SHAP values means these features make drug metabolized more by gut bacteria as shown in Figure 3 (C). Figures 6 and 7 in the Supporting Information show the summary of drug features contribution for other bacterial strains for activity and metabolism data models, respectively.

**Figure 3:**
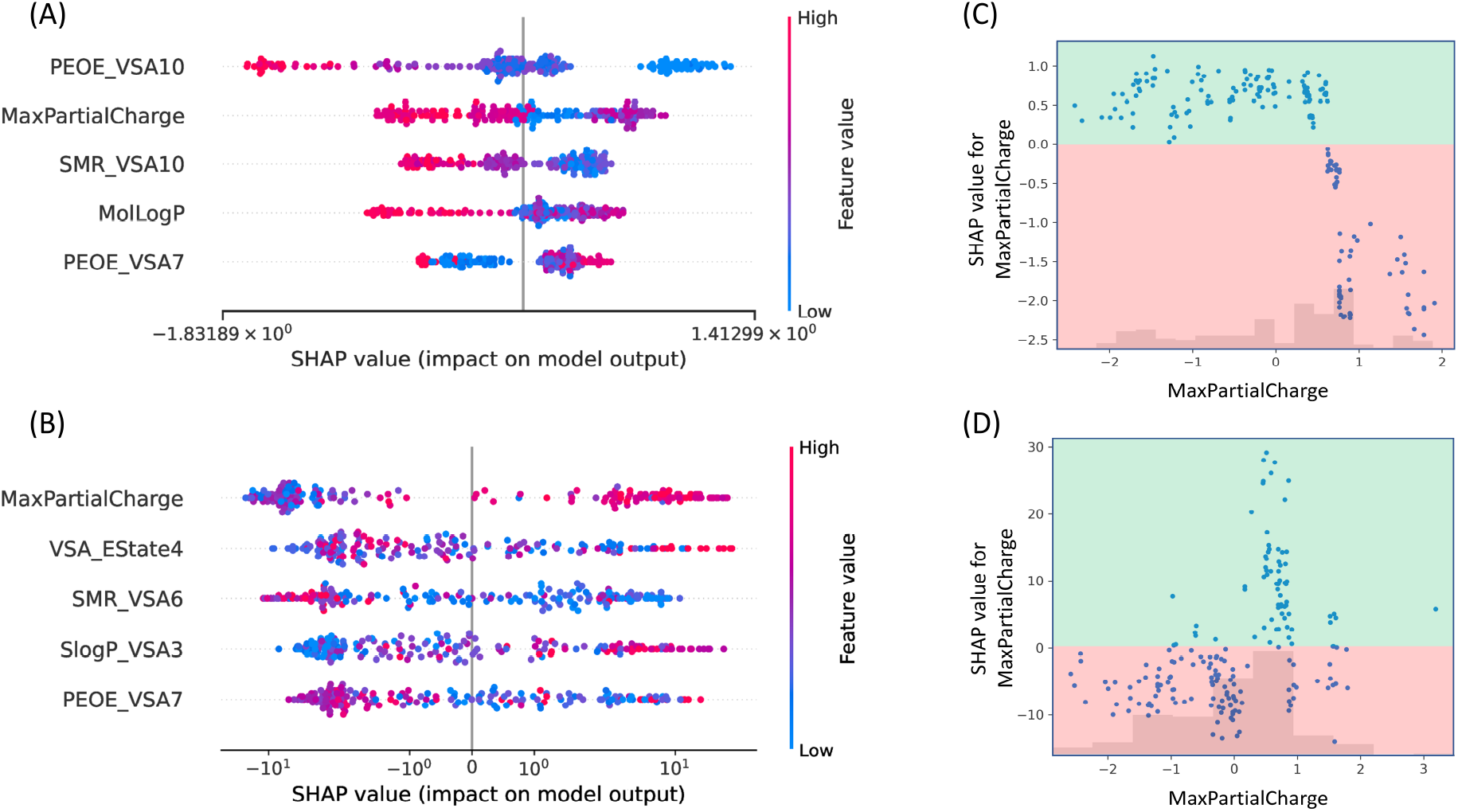
The contribution of key features in the model: (A) and (B) shows the summary of contributions of top key features in activity and metabolism data models. *MaxPartialCharge* is a common key feature in both data sets. Figure (C) and (D) shows the contribution of *MaxPartialCharge* in the activity and metabolism data models, respectively. The higher positive value of *MaxPartialCharge* gives negative SHAP values (y-axis), which means drugs with a higher positive charge will inhibit gut bacteria growth more (C). The higher negative value of *MaxPartialCharge* gives negative SHAP values (y-axis), which means drugs with a higher positive charge will get metabolized more by gut bacteria (D). Supporting Information Figures 6 and 7 shows the contribution of key features of the drugs in each bacterium model, and Figure 8 and 9 show the contribution of *MaxPartialCharge* in activity and metabolism data models, respectively.

### *MaxPartialCharge*: A common important feature

The *MaxPartialCharge* is the most common key feature of drugs in both the activity and metabolism data models, as shown in Figure 2 (C) and (D). It is the value of the maximum partial charge present on any of the atoms of the drug molecule. The contribution of *MaxPartialCharge* is different in both of the data models, as shown in Figure 3 (A) and (B) To further analyze the role of *MaxPartialCharge* feature of the drugs in detail, we used shap.plots.scatter function of the SHAP algorithm. A higher-positive value of *MaxPartialCharge* of the drug makes these drugs more harmful for the gut bacteria shown in Figure 3 (C). Previous studies show that cationic drugs can have antibacterial properties and effectively kill bacteria. Especially, the drug molecules that interact with the negatively charged bacterial membrane to kill bacteria.^39–41^ A more negative value of *MaxPartialCharge* of the drug makes these drugs more metabolized by gut bacteria shown in Figure 3 (D). Hence, this study shows that the chance of bacteria to metabolize drugs can increase with a higher negative partial charge present on the drug molecules, and a higher positive charge present on the drug molecules makes these drugs more harmful to the gut bacteria. Supporting Information Figures 8 and 9 show the role of the *MaxPartialCharge* of the drug in other models for different bacterial strains in activity and metabolism data, respectively.

### Prediction on FDA rejected drug

We used the available data from US-food and drug administration (FDA) website^42^ on rejected and accepted drugs (2019). We used our prediction models to predict the activity of FDA-rejected and FDA-accepted drugs on gut bacteria and activity of gut bacteria on FDA-rejected and FDA-accepted drugs. As shown in Figure 4 (A) and (B), the FDA-rejected drugs show more negative activity on gut bacteria, and get more metabolized by the gut bacteria, respectively. Although our study shows that FDA-rejected drugs have more negative gut-bacteria interactions, more detailed studies are required to conclude the activity of these drugs using physicochemical properties.

**Figure 4:**
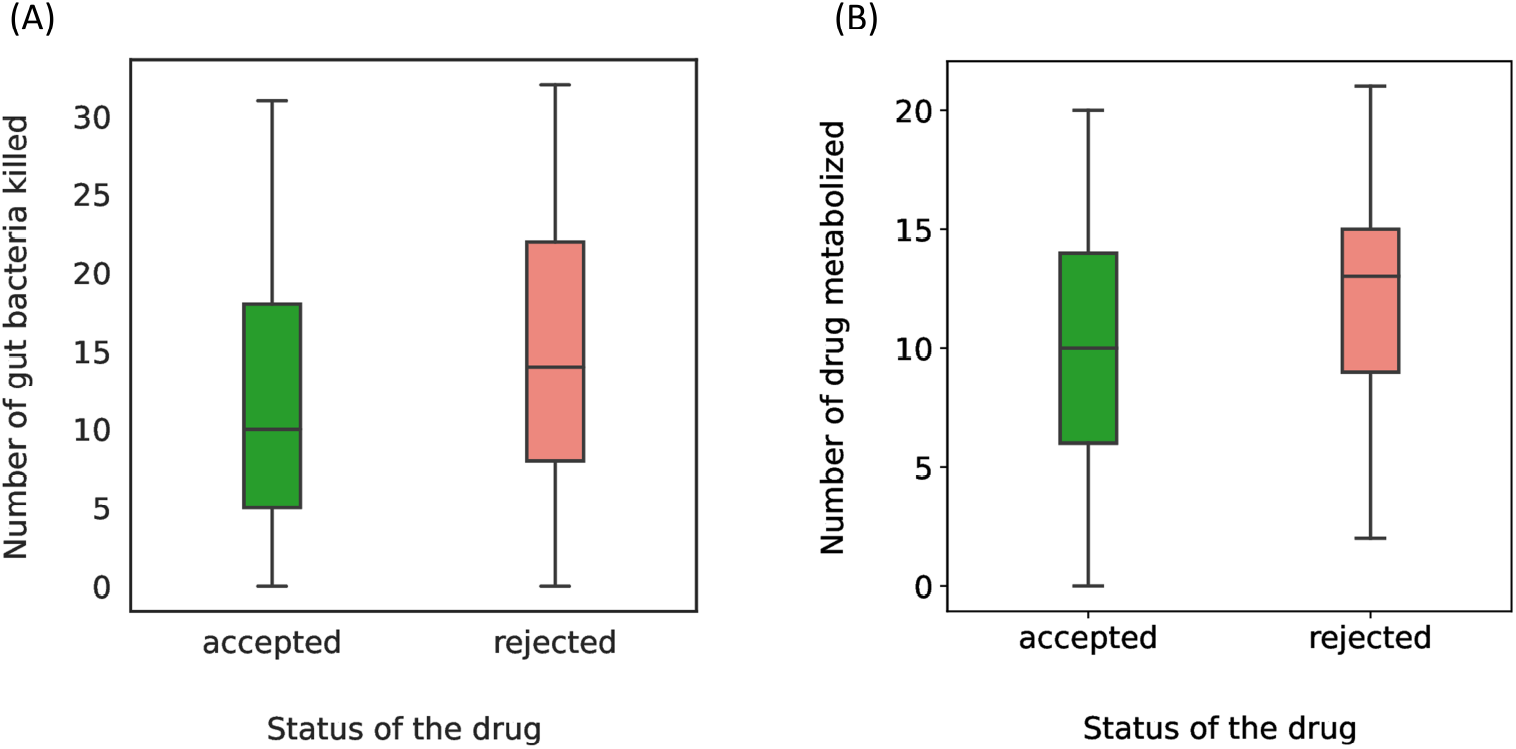
Predictions on FDA accepted and rejected drugs: We used both of the data models to predict the activity of drugs on gut bacteria (A) and metabolism by gut bacteria (B) on FDA-rejected and accepted drugs. The median of the box plot is higher for the prediction on the FDA-rejected drugs (red) than the prediction on the FDA-accepted drugs (green), as shown in (A) and (B), which means the predictions done by our model show that FDA-rejected drugs kill more gut bacteria and the rejected drug get metabolized more.

## Conclusion

The efficiency of drugs to the target decreases by the negative interactions of drugs and gut bacteria and can have side effects on the host’s health. In this work, we used interpretable machine learning models on drug-gut bacteria interactions to identify the crucial physico-chemical properties of the drug molecules leading to these interactions. We found that the drug features related to molecular weight, partial charge, and Van der Waal surface area play a key role in deciding the activity of the drug and gut bacteria on each other. One of the key findings of this study is that cationic drug molecules show negative activity on gut bacteria, and anionic drugs can get metabolized more by gut bacteria. While cationic drugs have shown promising antibacterial activity and anionic drugs get metabolized more, further research is needed to understand the drug-gut bacteria interactions since many other factors of the drug decide this activity. Machine learning models used to study the structure-property relationship of drug molecules are highly dependent on reliable input data. Input data sets used in this study are in vitro experimental data on an individual bacterium, neglecting the interactions between the bacteria. Since gut bacteria make a complex ecology, drug-gut bacteria interactions may differ in vivo conditions. For this study, we used 2D descriptors of the molecules so other available molecular descriptors could be used. If more data is available in the future, this model can give better results.

## Notes

The authors declare no competing financial interest.

## Additional Information

Data and codes are available at https://github.com/Himanshu0829/drug_gut_bacteria_interactions_ML-

## Acknowledgement

We thank Prof. Hemalatha Balaram and Prof. Sridhar Rajaram for extensive discussions on modeling and results. We thank Prof. Kavita Jain, Dr. Sruthi C. K., Dr. Malay Ranjan Biswal, Dr. Sudharshan Behra, Bidisha Bhatt, Lakshita Jindal, and Sayantan Maity for their helpful suggestions on writing the manuscript. H. J. wishes to express his sincere gratitude for the unwavering support that he has received from Prof. Kavita Jain throughout this project. H.J. also thanks CSIR for the fellowship.

## Notes

https://github.com/Himanshu0829/drug_gut_bacteria_interactions_ML-

### Competing Interest Statement

The authors have declared no competing interest.

