## Supporting Information for "Interpretation of Predictions in Drug-Gut Bacteria Interactions Using Machine Learning"

#### Input data

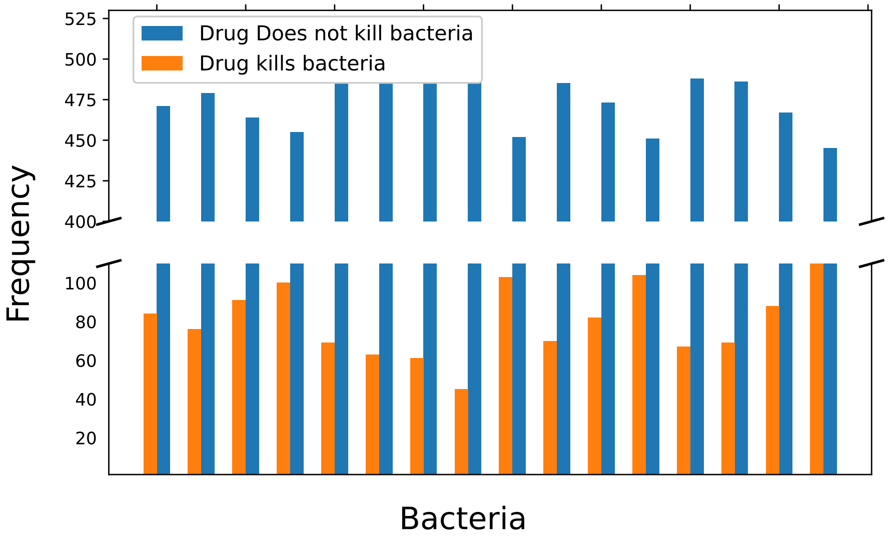


(A) Activity of drugs on gut bacteria data.

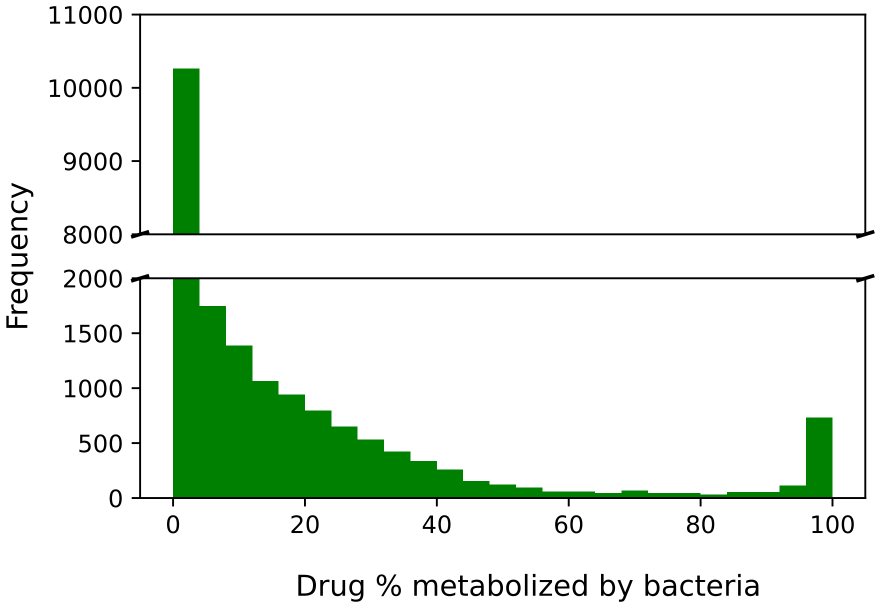


(B) Activity of gut bacteria on drugs data.

Figure 1: **Input Data:** (A) and (B) show that we have highly imbalanced input data. The activity data, the drugs that do not kill the gut bacteria, was downsampled to match the number of drugs that do not kill bacteria and make input activity data balanced (A). To make the metabolism data balanced, the drug which did not metabolize by gut bacteria was downsampled to make data balanced (B).

### Removed correlated drug features

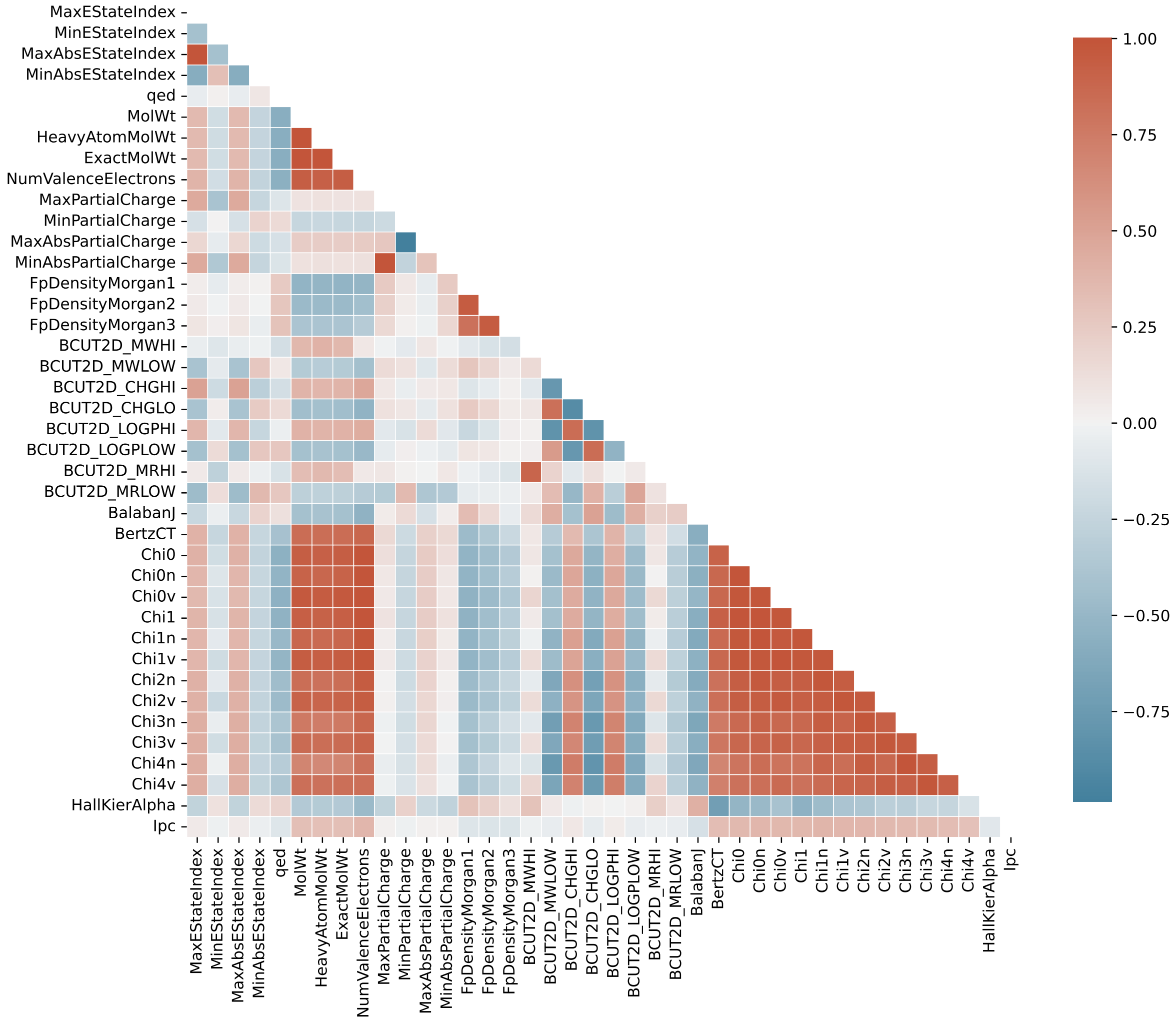


(A) The correlation between drug features and positive correlation is red, and the negative correlation is blue.

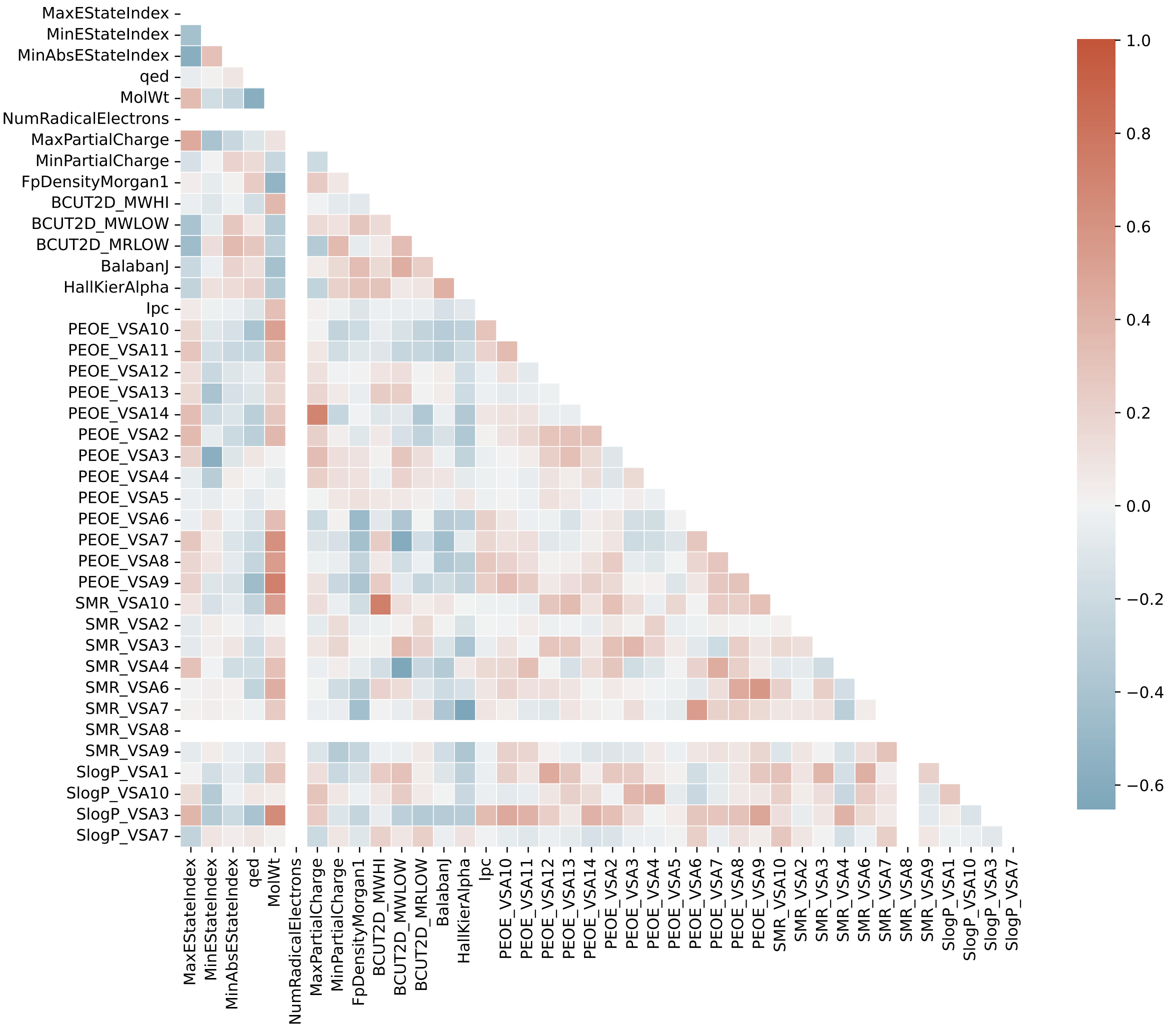


(B) Uncorrelated drug features.

Figure 2: **Feature Selection:** (A) shows that some drug features are highly correlated. So, we removed highly correlated features if the absolute Pearson correlation between any feature is more than 0.75 (B).

#### Quality and important drug features from first-generation prediction model

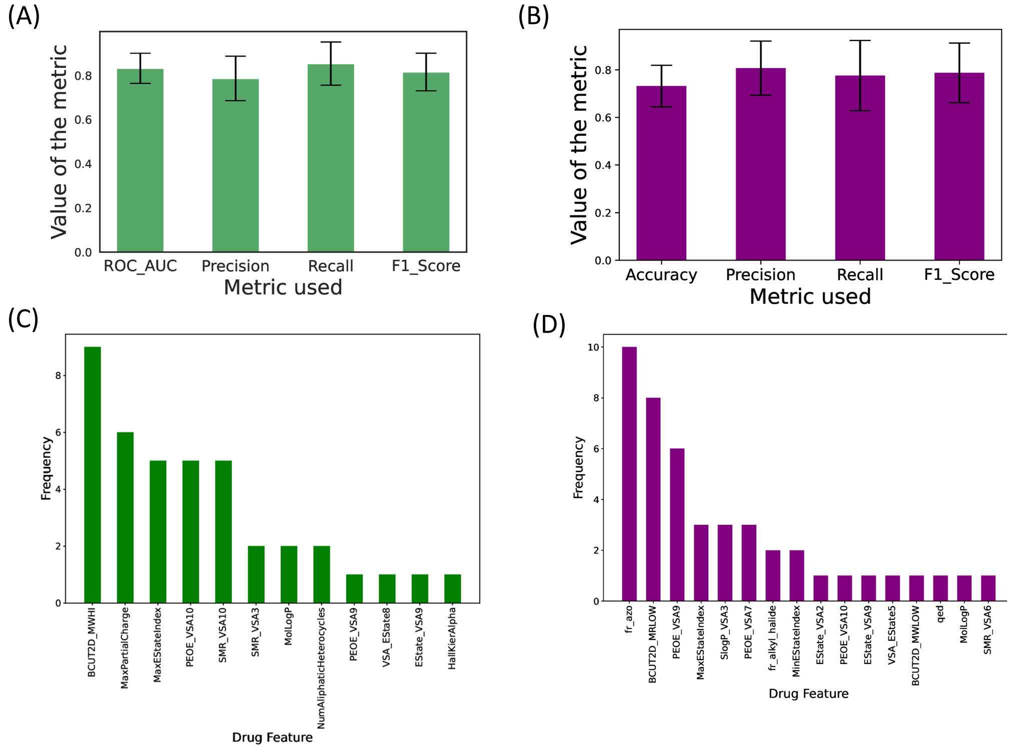


Figure 3: **Prediction quality and key features in first-generation models:** Figure (A) and (B) shows the mean and standard deviation of different metrics used to check the prediction qualities of the individual bacterium model for activity data and metabolism data, respectively. Using the SHAP algorithm, the top key drug physicochemical properties in the activity and prediction data models and plotted the frequency of the key features as shown in (C) and (D) in activity and metabolism data, respectively.

#### Quality of second-generation model

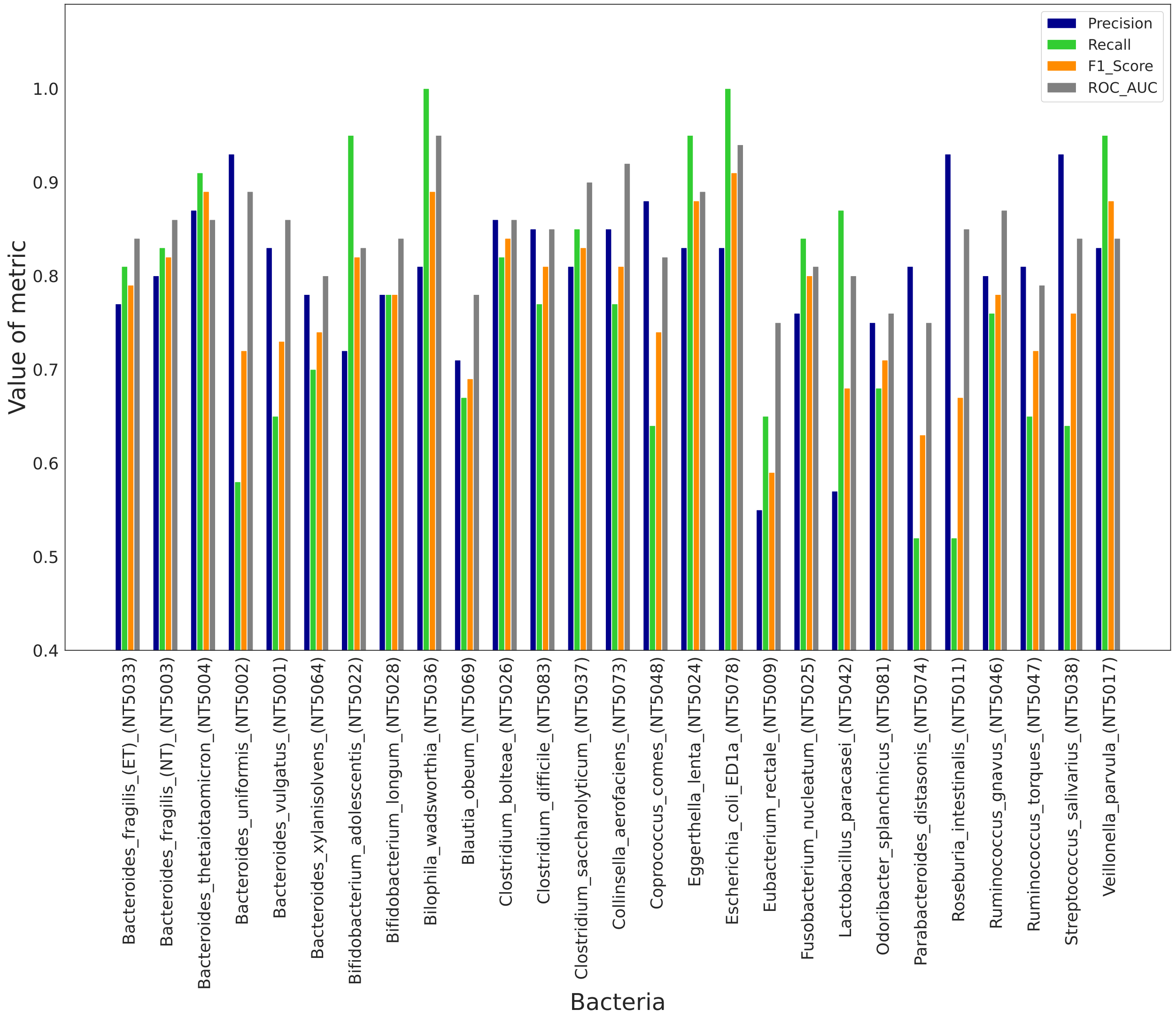


Figure 4: **The quality of the second-generation models for activity data:** We used AUROC, precision, recall, and F1score as the metric to check the quality of the prediction model. We selected those bacteria models with AUROC of more than 0.75 for further study.

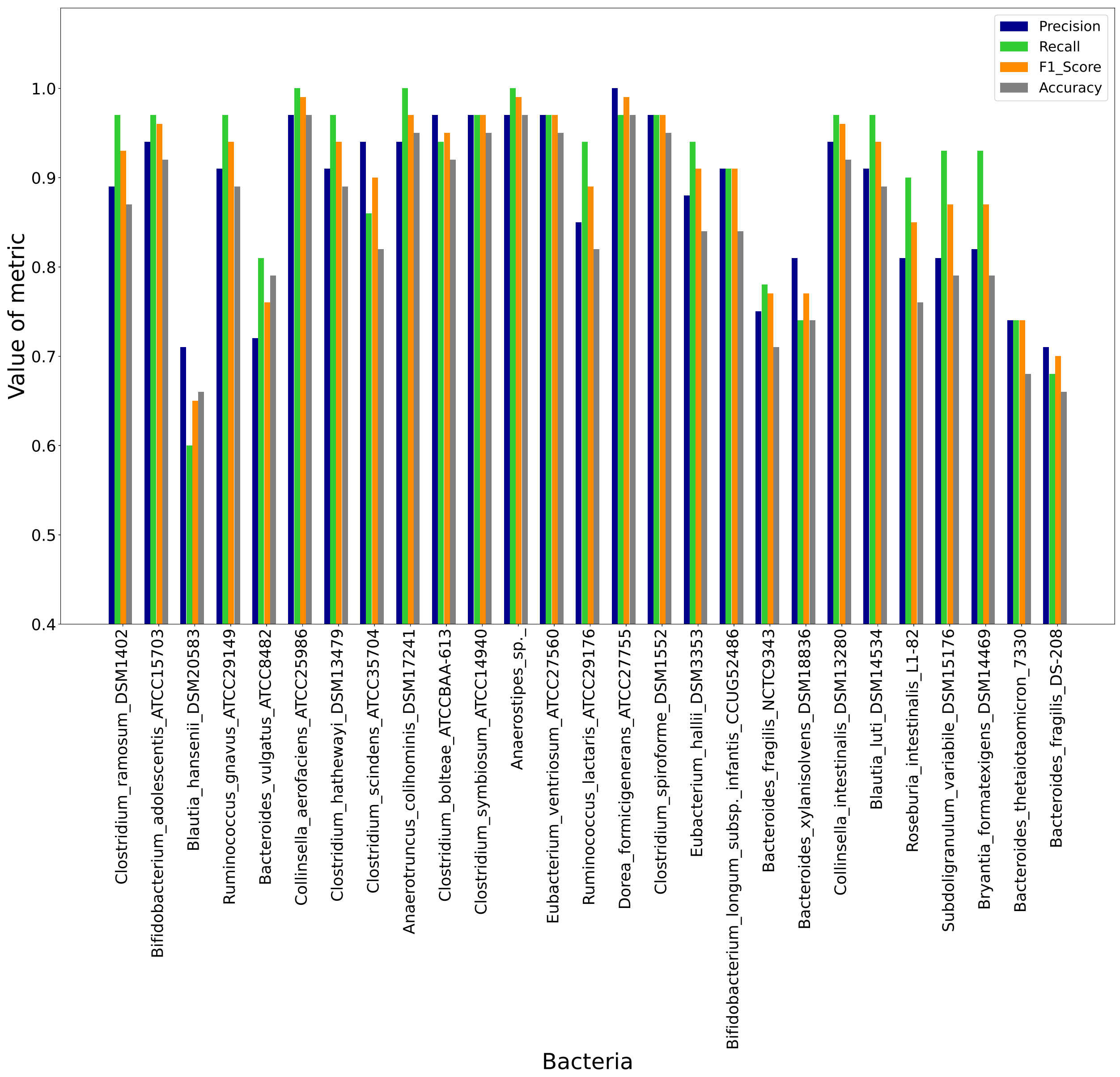


Figure 5: **The quality of the second-generation models for metabolism data:** We used accuracy, precision, recall, and F1score as the metric to check the quality of the prediction model. We selected those bacteria models with an accuracy of more than 0.70 for further study.

#### Definition of important features after second generation model

The 2D descriptors of the molecules are the numerical values of physicochemical properties. The definition and meaning of important molecular descriptors obtained from the 2 generation model is following;

1. **BCUT2D_MWHI:** This is an eigenvalue-based molecular descriptor calculated using an adjacent matrix. The adjacency matrix (M), of a chemical structure is defined by the elements $[M_{ij}]$ where $M_{ij}$ is one if atoms $i$ and $j$ are bonded and zero otherwise. The distance matrix (D) of a chemical structure is defined by the elements $[D_{ij}]$ where $D_{ij}$ is the length of the shortest path from atoms $i$ to $j$; zero is used if atoms $i$ and $j$ are not part of the same connected component. The BCUT descriptors are calculated from the eigenvalues of a modified adjacency matrix. Each $ij$ entry of the adjacency matrix takes the value $\frac{1}{\sqrt{(b_{ij})}}$ where $b_{ij}$ is the formal bond order between bonded atoms $i$ and $j$. The diagonal takes the value of the molecular weight. So, BCUT2D_MWHI means the higher eigenvalue of the matrix corresponding to higher atomic weight.[1]

2. **BCUT2D_MRLOW:**The BCUT descriptors are calculated from the eigenvalues of a modified adjacency matrix. Each $ij$ entry of the adjacency matrix takes the value $\frac{1}{\sqrt{(b_{ij})}}$ where $b_{ij}$ is the formal bond order between bonded atoms $i$ and $j$. The diagonal takes the value of the molecular refractivity of the molecules.[1]

3. **MaxPartialCharge:** Maximum partial charge present any of the atom in the molecule.[2]

4. **PEOE_VSA10:** The Partial Equalization of Orbital Electronegativities (PEOE) method of calculating atomic partial charges is a method in which charge is transferred between bonded atoms until equilibrium. Let $q_i$ denote the partial charge of atom $i$ as defined above. Let $v_i$ be the van der Waals surface area of atom $i$. PEOE_VSA10 is the Sum of $q_i$ where $v_i$ ranges from 0.10 to 0.15.[2]

5. **SMR_VSA10:** The Subdivided Surface Areas are descriptors based on an approximate accessible van der Waals surface area calculation for each atom, $v_i$ along with some other atomic property, $p_i$. A connection table approximation calculates the $v_i$. SMR_VSA10 is the sum of $v_i$ such that $R_i$ ranges from 0.10 to 0.15.

6. **SMR_VSA3:** SMR_VSA3 is the sum of Molar Refractivity such that the van der

Waal surface area of the atom is in the range of 1.82 to 2.24.[3]

7. **EStateIndex:** The EStateIndex descriptor is the combination of contributions of individual atomic and bond properties, each of which is assigned a specific integer value. The properties include the number of lone pairs, pi electrons, heavy atoms, and atomic electronegativity, among others. The resulting EStateIndex values can be used to compare different molecules' electronic properties and identify structural features that contribute to specific electronic effects.[3]

8. **quantitative estimation of drug-likeness (QED):** QED is the combination of different descriptors that characterize the molecular properties of the drug molecules, such as its molecular weight, log P, polarity, and solubility, which indicates its likelihood of success as a drug.[4]

9. **MlogP:** The MLogP value is calculated using a fragment-based approach, where the molecular structure is broken down into smaller fragments, and the contribution of each fragment to the overall lipophilicity of the molecule is estimated. The final MLogP value is obtained by summing up the contributions of all the fragments in the molecule.[5]

### Interpretation of predictions

#### Contribution of drug features to the models

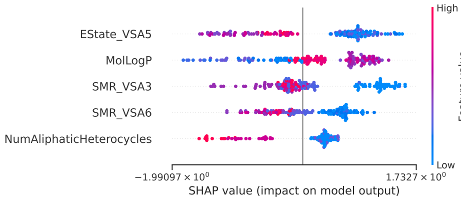


(A) Akkermansia_muciniphila_NT5021

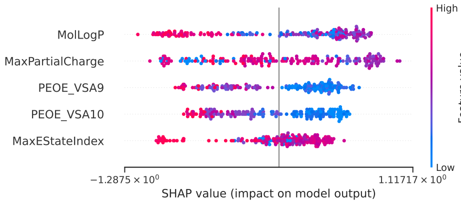


(B) Parabacteroides_distasonis_NT5074

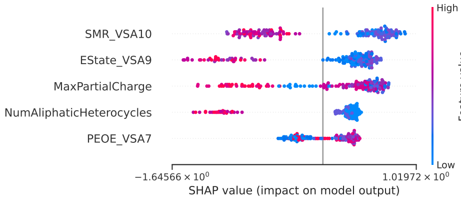


(C) Bifidobacterium_longum_NT5028

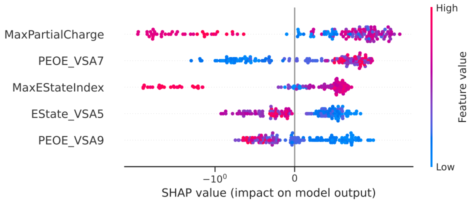


(D) Bilophila_wadsworthia_NT5036

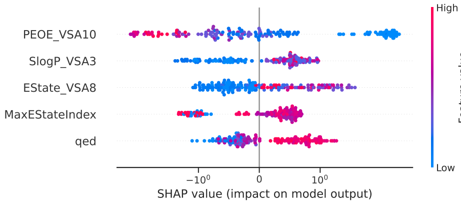


(E) Eggerthella_lenta_NT5024

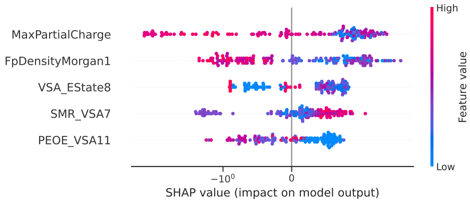


(F) Escherichia_coli_ED1a_NT5078

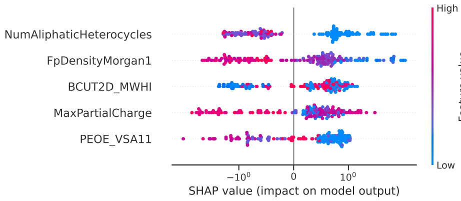


(G) Escherichia_coli_IAI1_NT5077

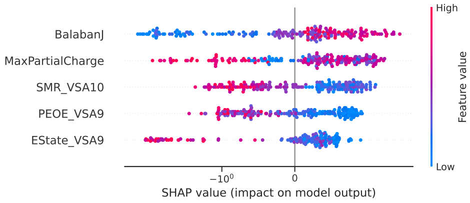


(H) Eubacterium_rectale_NT5009

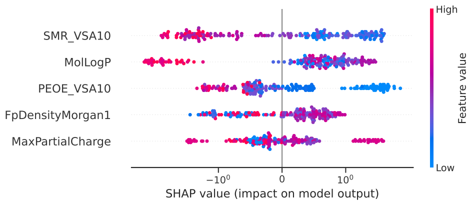


(I) Parabacteroides_merdae_NT5071

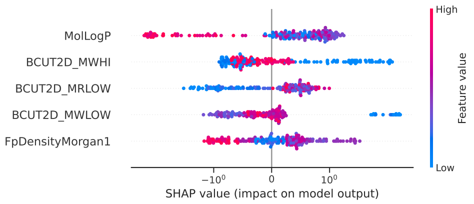


(J) Prevotella_copri_NT5019

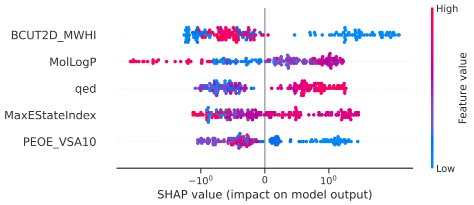


(K) Ruminococcus_gnavus_NT5046

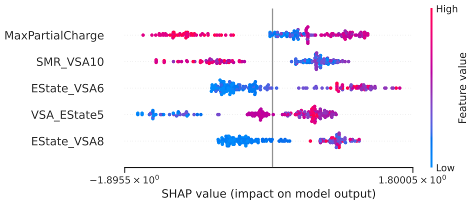


(L) Veillonella_parvula_NT5017

Figure 6: **Key features' contributions in the model:** Summary of contribution of all the top key features of the drugs in different models for individual bacterium models for activity data.

#### MaxPartialCharge: A common important feature

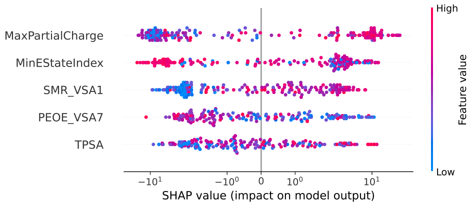


(A) Bacteroides_fragilis_DS-208

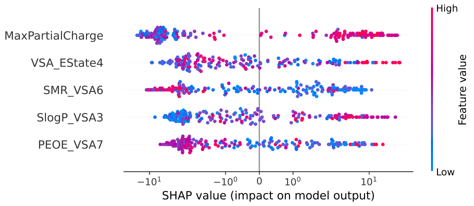


(B) Bacteroides_fragilis_NCTC9343

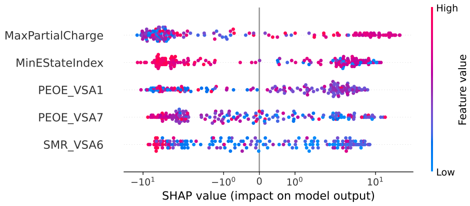


(C) Bacteroides_fragilis_TB9

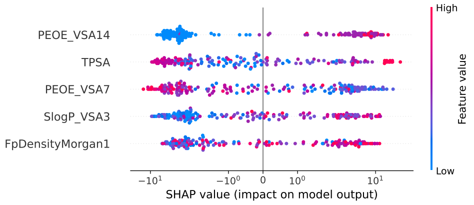


(D) Bacteroides_thetaiotaomicron_3731

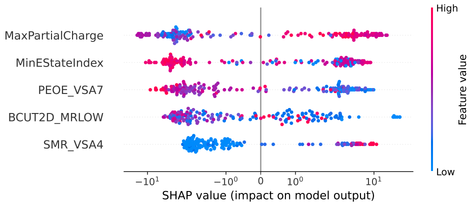


(E) Bacteroides_uniformis_ATCC8492

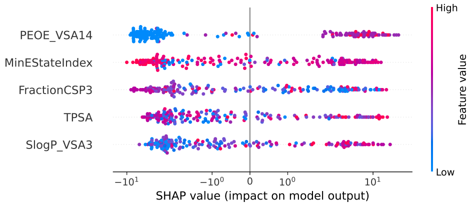


(F) shap_Bacteroides_xylanisolvens_DSM18836

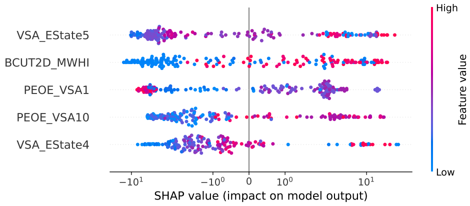


(G) Bifidobacterium_breve_DSM20213

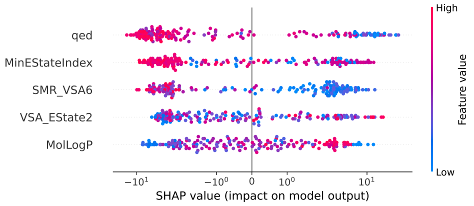


(H) Blautia_hansenii_DSM20583

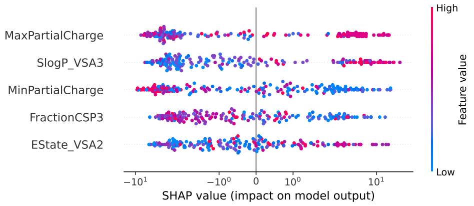


(I) Odoribacter_splanchnius

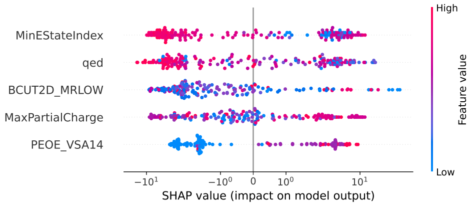


(J) Parabacteroides_distasonis_ATCC8503

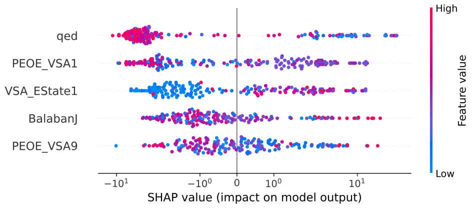


(K) Roseburia_intestinalis_L1-82

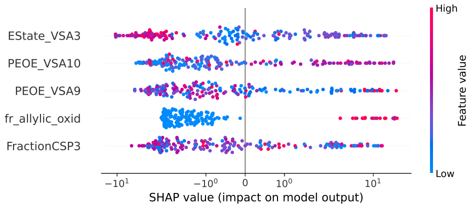


(L) shap_Ruminococcus_lactaris_ATCC29176

Figure 7: **Key features' contributions in the model:** Summary of contribution of all the top key features of the drugs in different models for individual bacterium models for metabolism data.

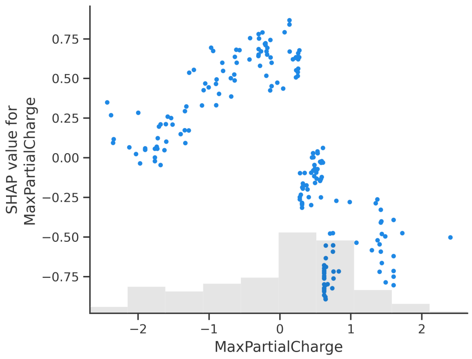


(A) Bacteroides_fragilis_(NT)

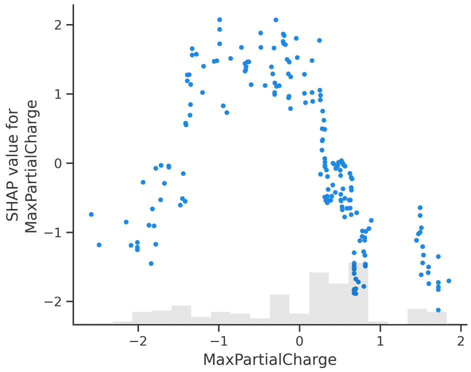


(B) Bacteroides_vulgatus

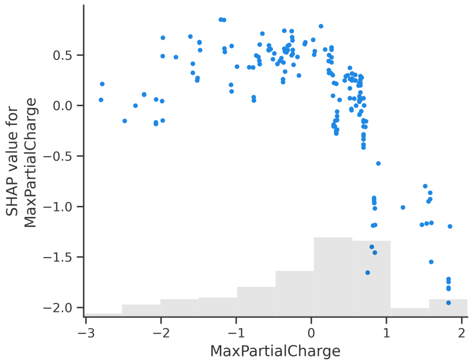


(C) Bifidobacterium_adolescentis

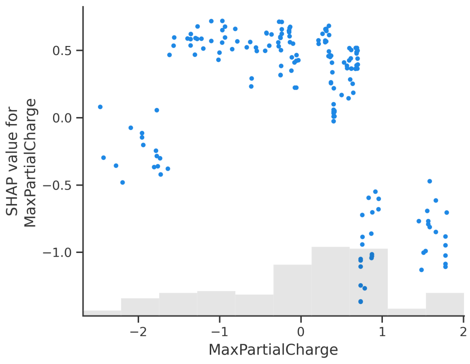


(D) Bifidobacterium_longum

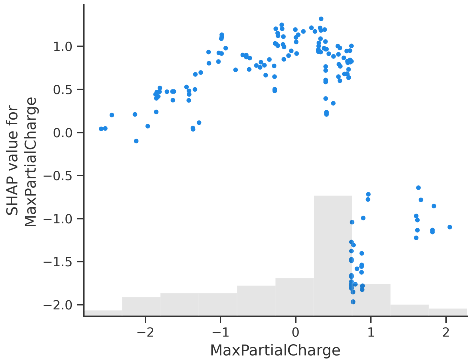


(E) Bilophila_wadsworthia

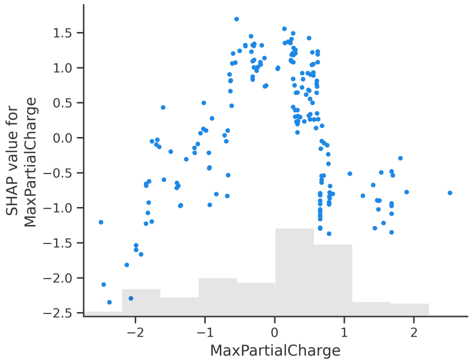


(F) Collinsella_aerofaciens

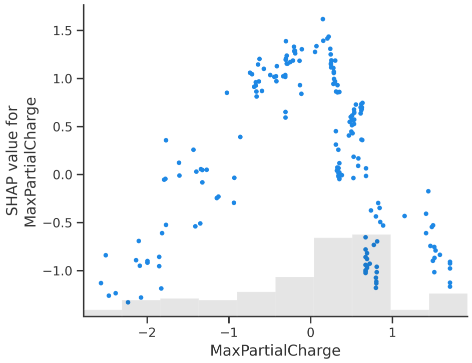


(G) Dorea_formicigenerans

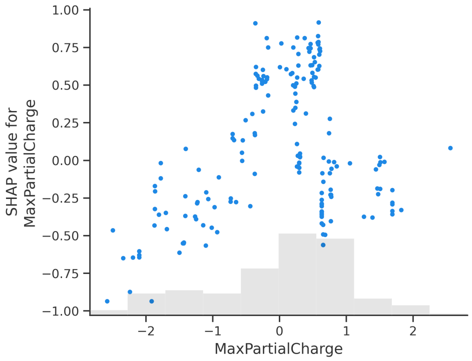


(H) Eggerthella_lenta

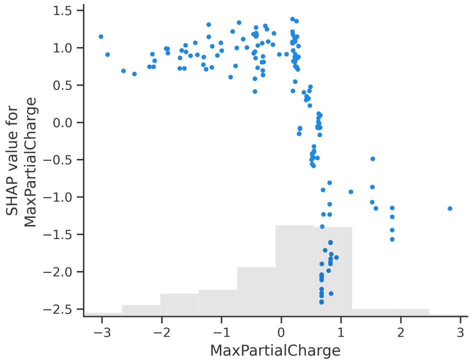


(I) Escherichia_coli_ED1a

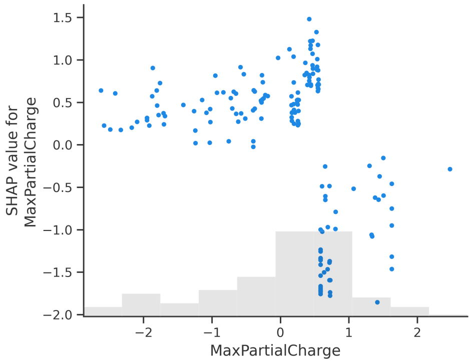


(J) Escherichia_coli_IAI1

Figure 8: **Contribution of *MaxPartialCharge* in the individual bacterium model in activity data.** A higher positive partial charge of the drug shows a negative value for SHAP value (y-axis), which means the drug will have more chance of metabolizing by gut bacteria. However, the higher negative partial charge on the drug shows positive values of SHAP, which means the drug has less chance of getting metabolized by gut bacteria.

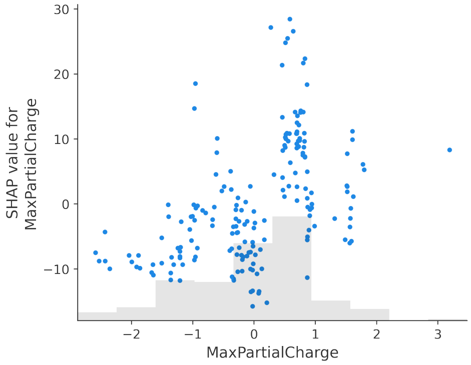


(A) Bacteroides_fragilis_DS

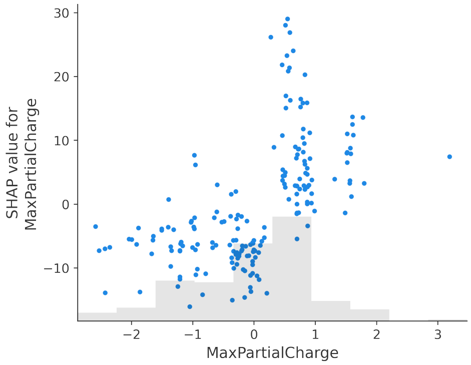


(B) Bacteroides_fragilis

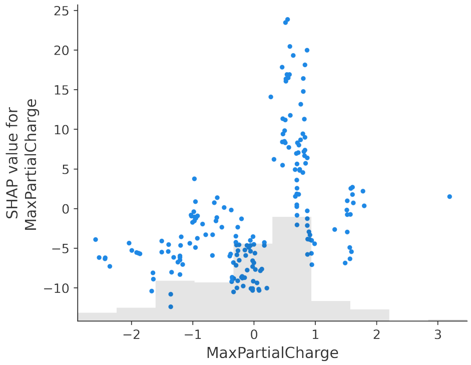


(C) Bacteroides_fragilis_T(B)

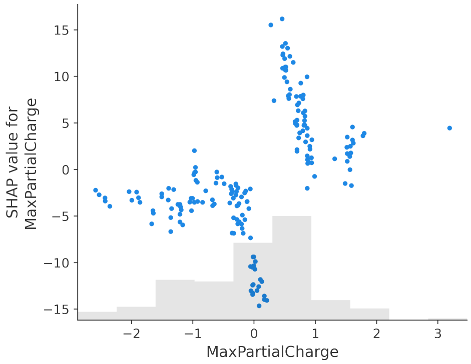


(D) Bacteroides_uniformis

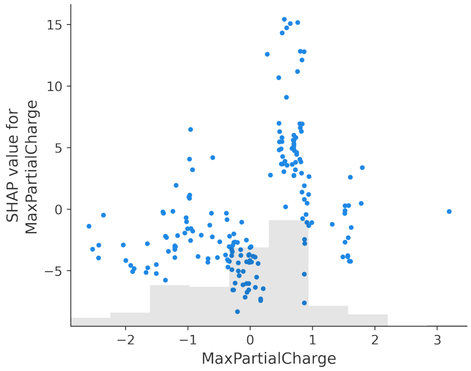


(E) Odoribacter_splanchnius

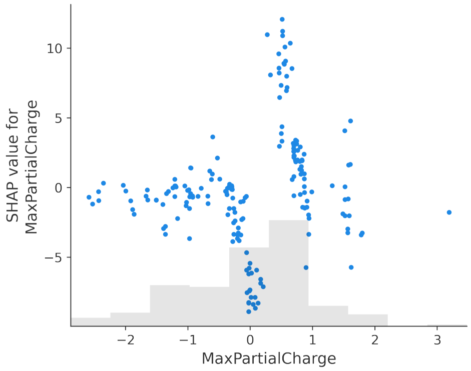


(F) Parabacteroides_distasonis_ATCC8503

Figure 9: **Contribution of *MaxPartialCharge* in the individual bacterium model in metabolism data.** A higher positive partial charge of the drug shows a positive value for SHAP value (y-axis), which means the drug will have less chance of metabolizing by gut bacteria. However, a higher negative partial charge drug shows negative values of SHAP, which means the drug has a higher chance of getting metabolized by gut bacteria.
